# Discrete mechanical growth model for plant tissue

**DOI:** 10.1101/459412

**Authors:** Louis D. Weise, Kirsten H. W. J. ten Tusscher

## Abstract

We present a discrete mechanical model to study plant development. The method is built up of mass points, springs and hinges mimicking the plant cell wall’s microstructure. To model plastic growth the resting lengths of springs are adjusted; when springs exceed a threshold length, new mass points, springs and hinges, are added. We formulate a stiffness tensor for the springs and hinges as a function of the fourth rank tensor of elasticity and the geometry of the mesh. This allows us to approximate the material law as a generalized orthotropic Hooke’s law, and control material properties during growth. The material properties of the model are illustrated in numerical simulations for finite strain and plastic growth. To solve the equations of motion of mass points we assume elastostatics and use Verlet integration. The method is demonstrated in simulations when anisotropic growth causes emergent residual strain fields in cell walls and a bending of tissue. The method can be used in multilevel models to study plant development, for example by coupling it to models for cytoskeletal, hormonal and gene regulatory processes.

## Introduction

Plant development is a complex process, it self-organizes using hormonal, gene regulatory and mechanical processes that act on multiple length- and time-scales and are linked via feedback loops. For instance, active and passive transport of hormones affects gene expression, but is also controlled by it: e.g. the hormone auxin [1] affects the expression of its own transport proteins [2], while also the expression of those transporters regulates the distribution of auxin [3]. Moreover, gene regulatory and hormonal processes govern the plant’s mechanical processes [4, 5], such as the cell wall’s expansive growth, rupture, and cell division. However, also mechanical determinants feed back on genetical and hormonal processes [6, 7]. For example, auxin guides root growth [8, 9], while a bending of the root also causes changes in auxin patterning [10, 11].

It is difficult to understand the consequences of such feedback loops from experiments alone. This is because experimental measurements are typically limited in their spatial and temporal scope, as well as in the number of processes that can be studied simultaneously. Mathematical modeling has been shown to be a valuable tool to study complex biological systems compassed of processes happening on different time- and length- scales and involving feedback loops [12, 13]. As a consequence, joint experimental and modeling approaches have led to important insights in plant development [8, 10, 14, 15].

It is important for mathematical models for plant growth to include elastic properties, because plants are sensing mechanical clues and are responding to them [16]. For instance, during cell division when a new cell wall is built, it is oriented such that it optimally resists tensile stress [17]. Also plant cells reinforce their wall by adding new microfibrils in the orientation of the highest stress [18]. To develop such models for plant growth mechanics is challenging, since plant tissue is a complex material that can be highly anisotropic and undergoes both elastic and plastic deformations during development. Plant cells are typically under a high turgor pressure, while being encased in stiff cell walls that resist this pressure [19]. Dynamical regulation (genetically and hormonally controlled) of the cell wall’s stiffness allows plastic growth [20], as well as the more extensive cell wall remodelling that is for example necessary for lateral root emergence [21].

The most established way to model the mechanics of plant development is in terms of continuum mechanics [22–26]. The application of continuous mechanical models has resulted in important insights, for example the demonstration that mechanical signals together with auxin patterning synergystically regulate plant shoot morphogenesis [15, 27]. A main advantage of the continuum mechanics approach is that material properties of the tissue are implicitly maintained during deformation, which is important since stress- and strain fields are relevant for plant morphogenesis [17, 27]. However, continuous methods have also theoretical and practical limitations. The plant’s cell wall is inherently discrete, consisting of networks of crosslinked fibrils. Mechanical processes on this scale, such as stiffening, loosening or rupture of fibrils are important for plant development (e.g. during lateral root formation), but are difficult to access locally with continuous models. Technically, it is a challenging problem to implement a continuum mechanics approach in a computationally efficient manner.

Discrete mechanical modeling offers an interesting alternative to continuum mechanics. A discrete mechanical model can mimic the microstructure of the plant’s cell wall, while being more easy to implement in a computationally efficient way. However, a main drawback of discrete mechanical models is that material properties are often not well defined: typically relations between discrete mass points are described, yet the stress-strain relation is not explicitly formulated [28]. Furthermore, deformations, e.g. due to growth, change the geometry of the mesh, thus causing undesired changes in material properties. In this paper we develop a discrete mechanical model to study plant development that is aimed at alleviating these limitations. We formulate a stiffness tensor for the mass point’s springs and hinges in terms of a generalized orthotropic Hooke’s law and the geometry of the mesh. Furthermore, we develop a remeshing method to control the mass point density and material properties during growth. Our model enables the incorporation of experimental data on elastic properties of plant cell walls. Finally, given the discrete nature of the model we can locally affect the stiffness of the model.

We demonstrate the model in simulations on anisotropic tissue growth. The model allows us to study strain fields and tissue bending that emerge due to anisotropic growth. The method can be coupled to existing models for hormone and gene regulatory networks and thus provides a valuable building block for multilevel models of plant development. The advantage of computational and numerical simplicity make our model an attractive method for researchers studying development of tissues involving growth mechanics of turgoid cells.

## Methods

### Main assumptions

First, we explain our main working assumptions: reduction of dimensionality, and usage of a simplified material law, before we explain the setting up of the model.

#### Reduction of dimensionality: plane stress assumption

Plant tissues are inherently three-dimensional. However, for many important research questions it is often reasonable to approximate plant tissue using simplified two-dimensional (2D) models. For instance, it has been shown in a 2D model that root bending may cause maxima in local auxin production [10]. Indeed, previous models for plant growth have used a 2D approximation, for instance the vertex- and hybrid vertex-midline models of Fozard et al. [29, 30] and Merks et al. [31]. A 2D approximation using the plane stress assumption is often made in shell models for plant tissue [27, 32]. The rationale behind this simplification comes from the observation that the cell walls of the outer cell layer (epidermis) of plant tissue is typically stiffer than its inner ones, and basically acts as a “tension-stressed skin” [33, 34]. Here we make use of the plane stress assumption to build a mechanical growth model for plant tissue. Plane stress occurs in thin plates where load forces act only parallel to them. We believe the plane stress assumption is most appropriate for applications on the level of the cell wall, which can be seen as a thin plate. However, to illustrate our method we will stretch the applicability of the assumption in two applications. In one application we consider anisotropic unidirectional growth in a slab of tissue which we connect to “root growth”. To model root growth the plane stress assumption is not well motivated (not a thin plate); one can argue that a rotational symmetry of the root allows a 2D approximation. We also illustrate the model in a setup in which asymmetric bidirectional growth happens in a thin sheet of tissue, similar to “leaf growth”. In this setup the we assume the leaf to be thin relative to the other two dimensions to justify the plane stress assumption.

#### Simplified material properties

Elastic properties of a material are formulated in terms of constitutive relations, equations that connect stress and strain. Constitutive relations of plant tissues are complex, as these tissues are typically anisotropic, and consist of distinct cell layers with divergent mechanical properties [35]. Furthermore, these properties are changing dynamically. For example, it has been shown that the stiffness of cell walls of *Arabidopsis thaliana* varies over one order of magnitude depending on the growth phase [36]. The material properties of plant tissue has been measured in experiments to be nonlinear using for example atomic force microscopy [37, 38]. However, we are not aiming on formulating a quantitative material relation, but want to capture only the main features to develop a method for qualitative applications. Therefore we use in our model a linear relationship between stress and strain and neglect higher order terms.

### Elastic model

Here we describe the setup of the discrete elastic model. We start with illustrating the mesh of the model which is built up by mass points, springs and hinges. For the springs and hinges we formulate a stiffness tensor in terms of the geometry of the mesh and the elasticity tensor. Then we explain how we describe elastic anisotropy and turgor pressure. Next we formulate an approximate material law. Finally we explain how we calculate the forces on mass points.

#### Mesh

We use a square lattice, where every mass point has four (if not at the border) neighboring mass points connected by springs (Fig 1A). A unit cell in this crystal lattice is shown in Fig 1B. In addition to springs to direct neighbors, there are hinges in each corner of a unit cell. The rationale for choosing this layout of mass points, springs and hinges is as follows. Plant tissues such as the root tip or hypocotyl are often anisotropic, and this anisotropy is caused by the presence of polarized cells. In these polarized cell types there are two principal cellulose fiber directions, one along the growth direction, and one perpendicular to it [39]. In contrast, some plant tissues are isotropic, containing apolar cells in which the cellulose fiber mesh is disoriented. Since we are interested in modeling plant tissues consisting of polarized cells, we choose a quadratic unit cell to mimic the two principal fiber directions of polar plant cells.

**Fig 1.**
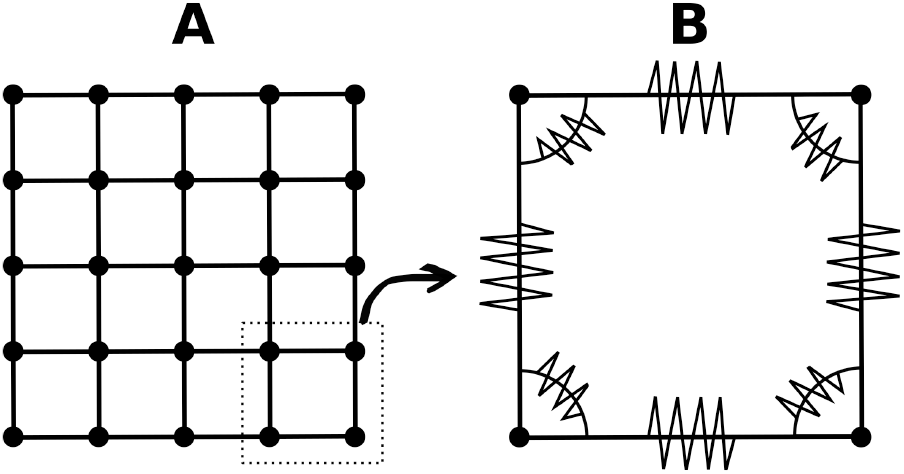
**(A) Mechanical mesh**. Dots indicate mass points. Springs are indicated as lines. Dotted contour indicates inset for subfigure. **(B) Unit cell**. Springs are indicated by zigzagging lines on straight lines. Zigzagging lines on curved lines connecting horizontal and vertical springs indicate hinges.

### Coupling to a continuous material law

The elastic properties of our model are determined by the geometry of the lattice unit cell and the stiffness of the springs and hinges. Here we will formulate these microscopic properties in the continuum limit from the macroscopic elastic properties of a linear elastic material.

The elastic energy density Ψ of a linear elastic material [40] is given by

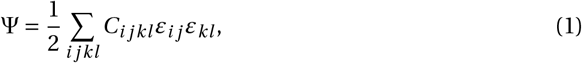

where *C*_*i jkl*_ are elements of the elasticity tensor, and *ε*_*i j*_ are components of the small strain tensor ***ϵ***. For an isotropic material the above simplifies to

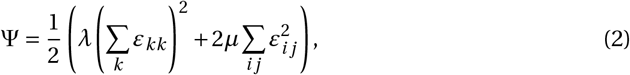

where *λ* and *μ* are the Lamé coefficients. The elements *σ*_*i j*_ of Cauchy’s stress tensor ***σ*** can be obtained (assuming constant temperature) by differentiating Ψ with respect to components from the strain tensor [41]

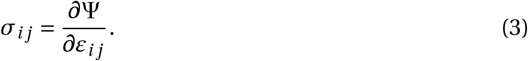

Substituting Ψ with Eq.2 we obtain the generalized Hooke’s law

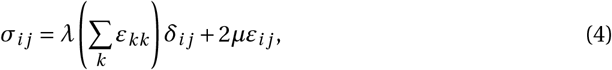

where *δ*_*i j*_ is the Kronecker delta.

To formulate a stiffness tensor for springs and hinges in terms of Lamé coefficients, we will take the following approach. First a description of the elastic energy density for the discrete model in terms of Lamé coefficients and the geometry of the mesh is found. Then we will do the same as above, find the elements of the stress tensor by differentiating the elastic energy density with respect to strain elements. Finally, we will compare the elastic energy density and stress tensor descriptions of our model to Eq.4 and Eq.1.

For the elastic potential of a spring in the x-direction we use (1/2)*k*(*y*_0_/*x*_0_)Δ*x*^2^, with spring stiffness *k*, and Δ*x* = *x* − *x*_0_ the change of length of a horizontal spring, where *x* is the actual length, and *x*_0_ is its resting length (similar terminology for the y-direction). Note that the term (*y*_0_/*x*_0_) takes the growth process into account: if the resting lengths in x- and y-direction are different while the relative deformation is identical in both directions, then the elastic potentials energy of both springs is identical in the unit cell. We use the potential *k*Δ*x*Δ*y* to account for the Poisson effect (similar to term 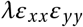 in Eq.2). Shear is described by means of four hinges per unit cell in terms of the potential (1/8)*κ*Δ*D*^2^ for each hinge, with *κ* the hinge stiffness, and Δ*D* the change of length of a diagonal in the unit cell. Fig 2 depicts a parallelogram which is used to formulate shear in terms of diagonals in a unit cell. Similarly to Eq.2 we write the elastic energy density 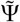 of the discrete mechanical model as

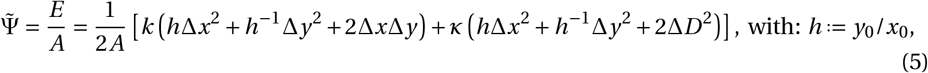

where *E* is the elastic energy of a unit cell, and *A* is its surface area. One horizontal spring is shared between two unit cells, whereas each unit cell contains two horizontal springs. Therefore, the net longitudinal strain of a single unit cell can be described as the deformation of a single spring *ε*_*xx*_ = Δ*x*/*x*_0_. Total shear of a unit cell is defined [40] as *τ* ≔ tan φ (compare Fig 2), and the components of the shear strain tensor (without growth) are *ε*_*xy*_ = *ε*_*yx*_ ≔ (1/2)*τ*. With growth (*x*_0_ ≠ *y*_0_) we find with same arguments as in [40]

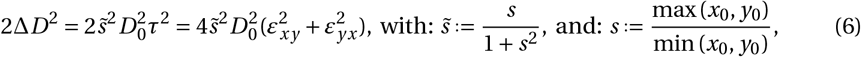

where 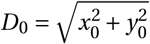 is the length of a diagonal of an undeformed unit cell. We rewrite 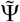 in terms of strain

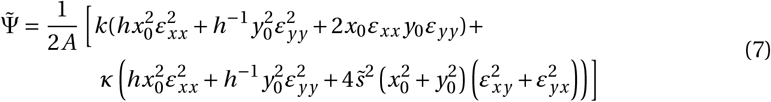

**Fig 2.**
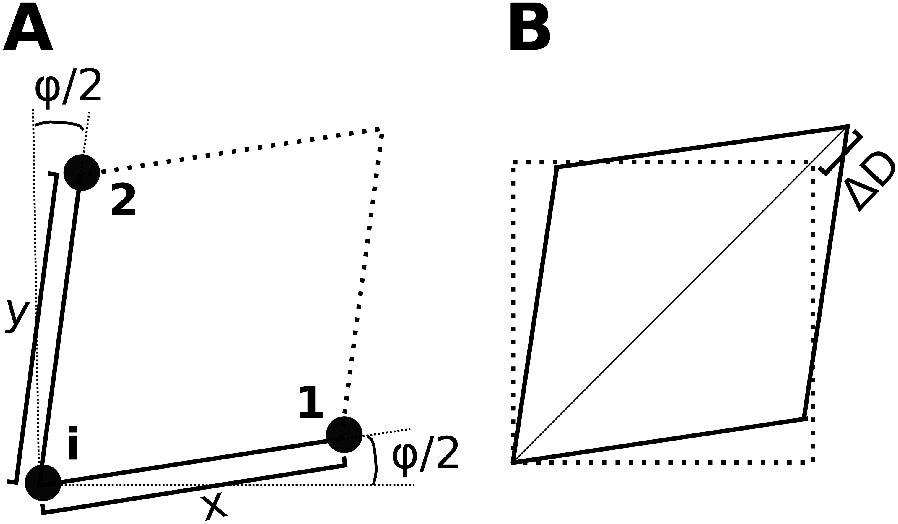
Illustration of parallelogram used to describe shear. **A Construction of parallelogram**. To calculate forces on point i, which results from one horizontal and vertical spring and one hinge, the positions of three mass points (***i*, 1, 2**) are used. We construct a parallelogram from these points (note thick dotted lines) to compute shear. **B Shear affects the length of the diagonals**. One diagonal of the parallelogram shortens, while the other lengthens by Δ*D*.

To get the elements of the stress tensor we differentiate 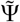 with respect to strain. We approximate

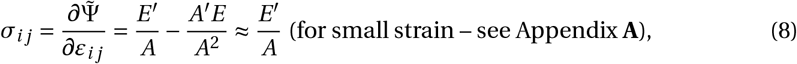

and compare the resulting expressions with Eqs.4 and 1 to get expressions for *k* and *κ*

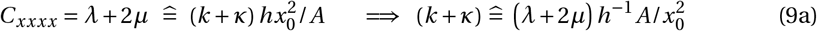

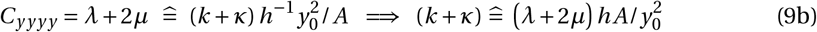

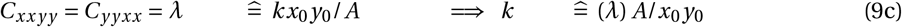

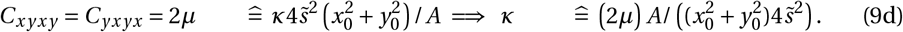

The stifness tensor of the discrete mechanical model 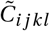 can now be written in terms of Lamé coefficients and geometric properties of an unit cell (we replace the terms with *k* and *κ*)

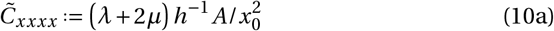

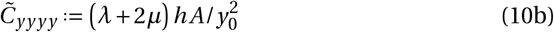

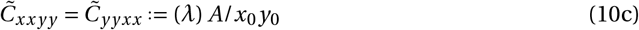

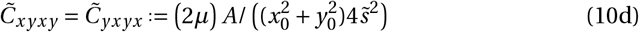

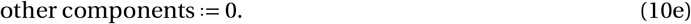

Finally, 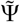 can be expressed similarly to Eq.1 as

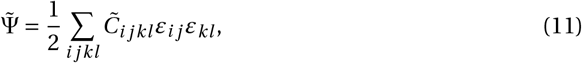

#### Anisotropy

In many plant tissues, cells are mechanically anisotropic. This is partly because of tissue specific polarized orientation of cellulose microfibrils in their cell walls [35]. The plant’s control of elastic cell wall properties plays a crucial part in plant morphogenesis. To model it generic and detailed bio-mechanical and chemical models have been developed [42, 43]. However, in this paper we do not model these physiologically important feedback loops, but account for anisotropy by defining two Young’s moduli, one for the x-direction *Y*_*x*_, and one for the y-direction *Y*_*y*_. Under plane stress Lamé coefficients are connected to Young’s moduli and Poisson’s ratio *v* [44] via

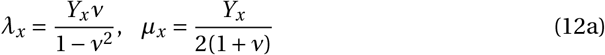

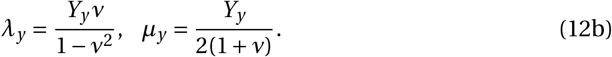

We define the shear modulus *μ* as the mean of the shear moduli of the isotropic materials characterized by either Young’s modulus (*Y*_*x*_, *v*) and (*Y*_*y*_, *v*)

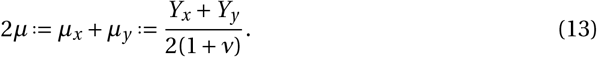

We rewrite the stifness tensor 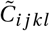 (Equation 10) for the anisotropic model

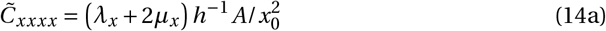

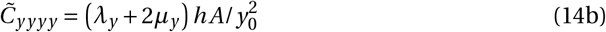

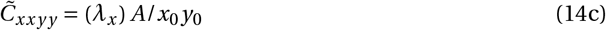

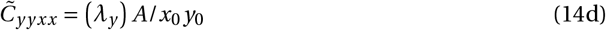

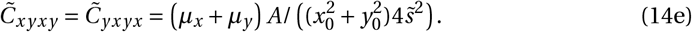

#### Constitutive relations

In the derivations presented above the small strain tensor *ϵ* was used to derive the properties of the springs and hinges. However, the small strain tensor is not suitable for finite deformations, because it is invariant to rigid body rotations [45]. Note, that the small strain tensor is related to the commonly used finite strain tensor [47]; in continuum mechanics we would get it by applying a polar decomposition on the small strain, to cancel out rigid body rotations. However, in our case, we do this implicitly, because we define strain in terms of relative length changes of springs, and angles between them. Thus, we can approximate the constitutive relations of our model in terms of the Biot strain tensor ***e*** [46].

With this we can approximate the elastic constitutive material relations for our anisotropic model as a generalized orthotropic Hooke’s law

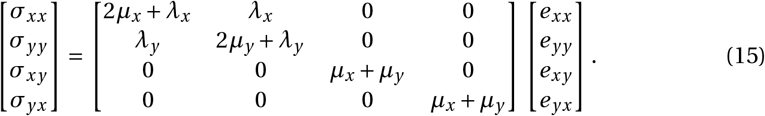

#### Turgor pressure

Assuming that turgor pressure *p*_*t*_ is constant across the tissue, i.e. all cells are equally turgid, the hydrostatic potential *E*_*hp*_ caused by *p*_*t*_ is

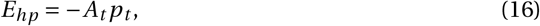

with *A*_*t*_ being the surface area of the tissue.

### Calculation of forces

In the following sections the calculation of the forces acting on the mass points will be explained. The forces will be used to compute the motion of the mass points.

#### Elastic forces

Elastic force on a mass point *i* (compare Fig 2) 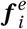 is the negative gradient of the elastic energy *E* with respect to ***i*** (note: *i* is an index, ***i*** is position vector of mass point *i*)

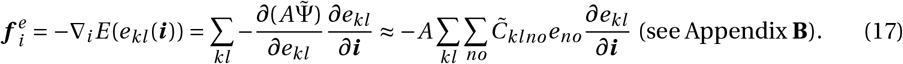

We expand it to (note that *e*_*xy*_ = *e*_*yx*_ = (1/2)*τ*)

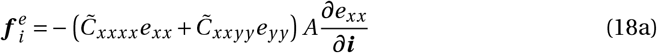

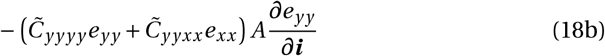

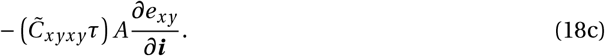

To solve Eq.18 we need to specify *A* and *e*_*i j*_ with respect to an individual mass point *i*, and get the derivatives *∂e*_*i j*_ /*∂****i***. Let us start with terms 18a,18b.

As can be seen from Fig 1A, a mass point inside the mesh is connected to four neighboring mass points and thus part of four unit cells (compare Fig 1B). A mass point at the boundary is part of two unit cells, and a point in the corner of only one unit cell. Therefore the mean surface area of the unit squares connected to the mass point *i* is used as variable *A* in terms 18a and 18b. These terms are calculated for all springs connected to mass point i. Thereby *e*_*xx*_ is the strain of one respective spring (similar in the y-direction). To calculate the force due to the Poisson effect, we use the strain in the “other direction”, e.g. *e*_*yy*_ for a spring in x direction (see term 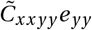), the mean *e*_*yy*_ strain of the adjacent springs in y-direction. Accordingly, we use the mean resting lengths of the adjacent springs in the y-direction to compute *y*_0_ in 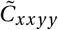. The derivatives of the direct strain elements with respect to coordinates of ***i*** are

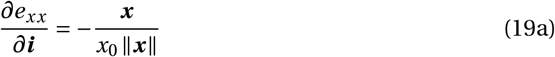

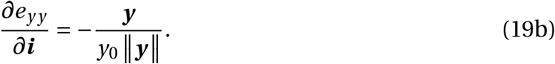

The term 18c is computed as follows:

the total shear force at a single mass point is described by its *N* adjacent hinges (*N* = 4 for a mass point in the medium, *N* = 2 for a mass point at the boundary of the mesh, and *N* = 1 for a point in the corner of the mesh). We define total shear strain at a single mass point *τ* as the mean of it’s *N* adjacent “hinge shear strains”

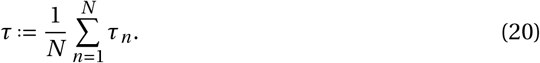

We use Fig 2 to illustrate how we calculate hinge shear strain *τ*_*n*_ for a hinge *n*. We use the springs in the x- and y-direction (see Fig 2) to define vectors: ***x***_*n*_ ≔ **1** − ***i***, ***y***_*n*_ ≔ **2** − ***i***, and use them to define *τ*_*n*_

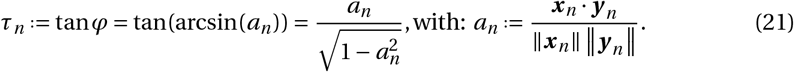

To calculate forces due to hinge shear strains in term 18c, we use the properties of the respective hinges: for the surface area of a unit cell we use *A*_*n*_ ≔ ||***x***_*n*_ × ***y***_*n*_|| (compare surface area of parallelogram in Fig 2). We also use respective springs in x- and y-direction of each hinge, to replace 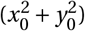 and 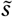 in Equation 10d with the corresponding hinge properties 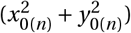 and 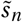. With this we write (see Appendix **C**) the shear force acting at a mass point *i* (term 18c) as a sum over the adjacent hinges

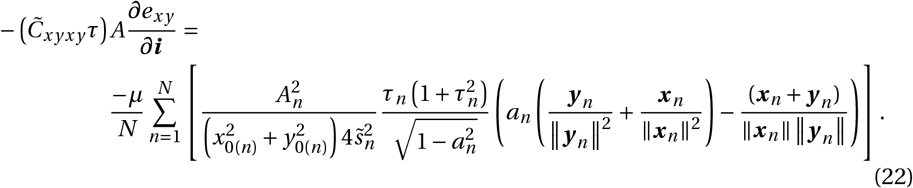

#### Forces due to turgor pressure

*A*_*t*_ in Eq. 16 is a planar non-self-intersecting polygon with vertices described by the position of mass points at the border of the tissue (*x*_1_, *y*_1_),…,(*x*_*n*_, *y*_*n*_) (vertices listed counterclockwise), thus *A*_*t*_ is given [48] by

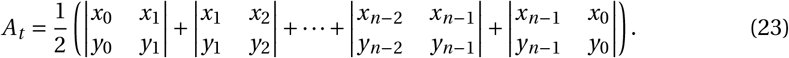

The “turgor force” 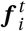 acting at a mass point *i* (if at the border of the medium), due to *p*_*t*_ is the negative gradient of the hydrostatic potential *E*_*hp*_ with respect to ***i***

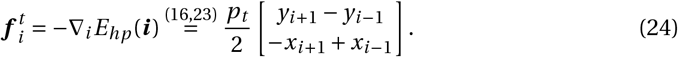

#### Viscous forces

We use a viscous force acting on every mass point to find the equilibrium configuration of the mesh

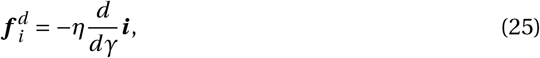

where *η* is the damping constant, and *γ* is the dimensionless integration time of the elasticity part of the model (see following section “Elastostatics” and “Numerical Methods”).

### Elastostatics

The motion of the mass points is described by Newton’s law of motion

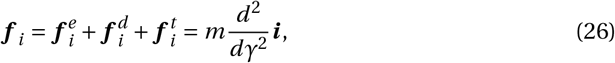

where parameter *m* is the mass of a mass point.

We make the common assumption [27, 32] that deformations happen at mechanical equilibrium. This can be understood from the fact that the plant’s growth processes are much slower than its elastic response to external forces. By solving Eq. 26 until mechanical equilibrium (***f*** _*i*_ = 0) we find the steady state configuration of the lattice. Note that the mass of a mass point *m* and the viscosity *η* have no physical relevance, because they do not affect the equilibrium configuration of the mesh. These parameters only fulfill numerical roles (convergence rate and precision).

### Plastic growth

Irreversible plant growth arises through the elongation of cell-walls of individual cells. This process involves cell wall loosening, expansion of the cell, and addition of new cell wall material restoring original cell wall stiffness [49]. During growth parts of the tissue are stretched or compressed to ensure a continuous tissue. Therefore, if an anisotropy in growth rates of cells is present in the medium residual stress and strain is build up in the tissue [50].

In the present paper we will apply our model to study basic effects of anisotropic growth such as emergence of residual strains and tissue bending. To model growth the resting length of springs is adjusted

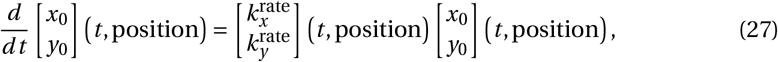

where *t* is the simulation time, and 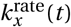 is the growth rate of a spring in the *x*-direction (similar terminology for the *y*-direction). This approach is similar to an evolving metric in the material manifold [51], and has been used before continuous mechanical growth models for plant tissue [32]. Note that the rate of change of the resting configuration is a complex function, Boudon et al. for instance formulated a strain-driven growth rate tensor [52]. Here we simply impose such growth functions to illustrate the elastic properties of our model during asymmetric growth. Note that this approach allows growth in two principal directions; however, can not describe a plastic shear growth.

#### Remeshing

The mass point density is affected by the growth process. Therefore, when springs exceed a threshold length of 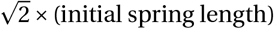, we add new mass points, springs and hinges to mimic the deposition of new cell wall material accompanying the later stages of cell expansion. We do this via the algorithm depicted in Fig 3. This figure shows that “loose mass points” can emerge in the medium during growth, points that have only three instead of four neighbors. We will see later in the results section that such loose points emerge when anisotropic growth causes local remeshing. However, we still calculate four hinge strains (see Eq.21) for such a loose point. For this an auxiliary point (see Fig 4) is assumed (on which no forces act). We calculate vector ***y*** using the loose mass point’s coordinates and the coordinates of the auxiliary point (similar for ***x*** when a loose end is pointing sideways). As resting distance *y*_0_ the resting distance of the left neighbor is used (similar, when a loose end is pointing sideways *x*_0_ of the upper neighbor is used).

**Fig 3.**
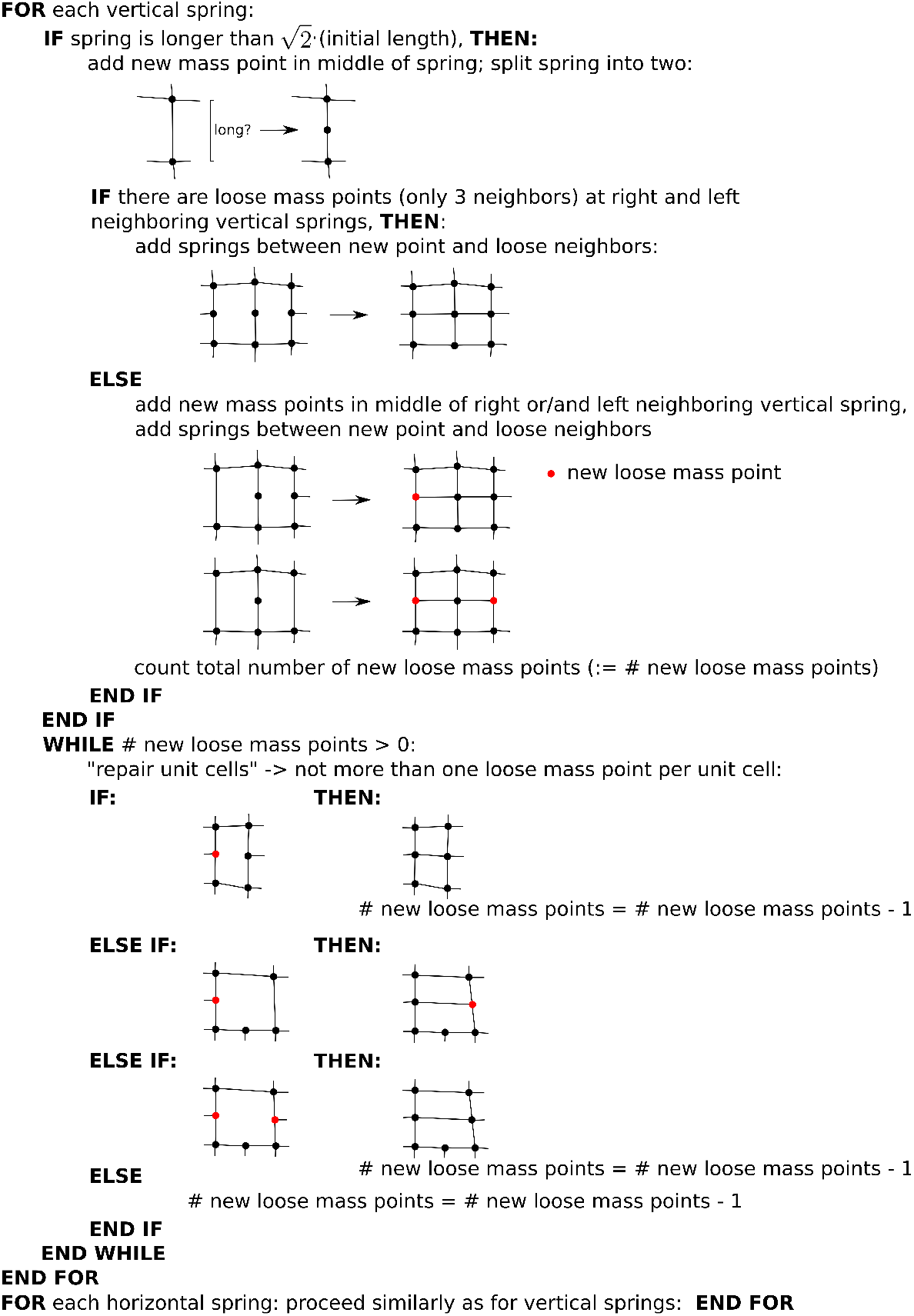
Illustration of remeshing algorithm. Dots: mass points, lines: springs.

**Fig 4.**
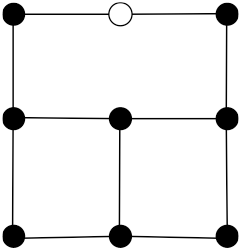
Illustration of usage of auxiliary point to calculate hinge strain for a loose mass point. The auxiliary point (white dot) is located in the middle of a spring between next-next neighbors Filled dots: mass points, black lines: springs.

### Numerical methods

We solved the equations of our model by combining explicit Euler integration for the growth equation (Eq. 27), and Verlet integration [53] to solve the equations for the motion of the mass points (Eqs. 26).

The position of a mass point ***i*** at integration time *γ* + *hγ* is computed with

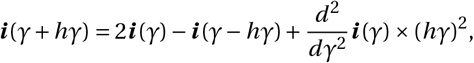

where *hγ* = 0.01 is the dimensionless Verlet integration time step and *γ* is the integration time. For the initial time step

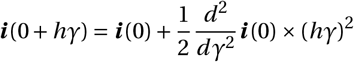

is used. The acceleration of a mass point *i* is found for each time step with

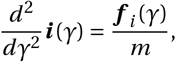

where *m* = 1*μg* is the mass of a mass point. The velocity of a mass point (to calculate the viscous force) in Eq.25 is computed with

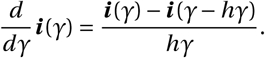

For Euler integration of the growth equation (Eq.27) we used an integration time step of *ht* = 1 *min*. The resting distances of springs connected to a mass point after a growth step are computed with

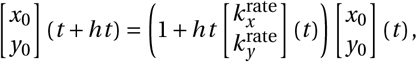

where *t* is the simulation time in minutes.

We solve the model as follows: from each time integration step of the growth model (Eq.27) a new set of resting distances of springs is obtained, which is passed on to the elasticity part of the model, where forces are computed (Eqs. 18,24,25), and the equations for the motion of the mass points (Eqs. 26) are solved, until the sum of forces on every mass point is below the convergence threshold *thr* = 0.05 *μN*.

In this study we used model setups of initially rectangular meshes of various lengths. As initial resting lengths of springs 1 *μm* was used. To compute viscous forces (Eq. 25) *η* = 1 *N* × *hγ*/*μm* was used as a damping constant. Several boundary conditions were used. For simulations shown in the results section a free floating medium (no degree of freedom is restrained, all mass points move freely) is used. In this setup a rigid body translation and rotation can occur; however, we perform a linear transformation to cancel these out rigid body motions (model is in moving frame of reference). In addition, in simulations to characterize the material properties of the model we either fix all boundaries of the model, or restrain points on specific boundaries to move on a line.

We set the parameter values of the model to connect it to experimental data on the model plant *Arabidopsis thaliana*. In experiments on roots, in which a rapid local growth was induced, a maximal relative elemental growth rate of one third per hour has been measured [54]. This corresponds to a maximal growth rate 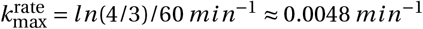, which we use as an upper boundary. In addition, we used as material properties Young’s moduli in the range of [20; 80] *MPa* · *m*, similar to reported experimental data on cell walls [55] and previous modeling work [27]. We varied turgor pressure in a range of [0; 1] *MPa* · *m*, corresponding to values measured on the root [56]. There are to our knowledge no precise measurements of the Poisson ratio *v* of the primary cell wall. However, it is established that the primary plant cell wall is a partially compressible material (*v* < 0.5) [57]. Previous modeling work [27] assumed *v* = 0.2. We vary *v* in a range of [0.1; 0.5] for simulations characterizing the model’s material properties, and use *v* = 0.2 for the illustration of the full model (Figs 10,11).

### Model parameter validation

The numerical coupling and integration of the Euler and the Verlet scheme require the choice of several parameters. In this section we explain our parameter choices to assure efficient and stable computations.

#### Integration Parameters

For the integration of the growth model (Eq.27) we use an Euler integration step *ht* = 1 *min*. We performed test simulations in which we used halfed and quartered values for our choice of *ht*. We found that our choice yields consistent results (see Fig. 5). For solving the motion of the mass points Eqs.26 we use a Verlet integration time step of *hγ* = 0.01 (as in [58, 59]). Thus, this setting allows efficient and stable computations of new configurations of the mechanical mesh.

**Fig 5.**
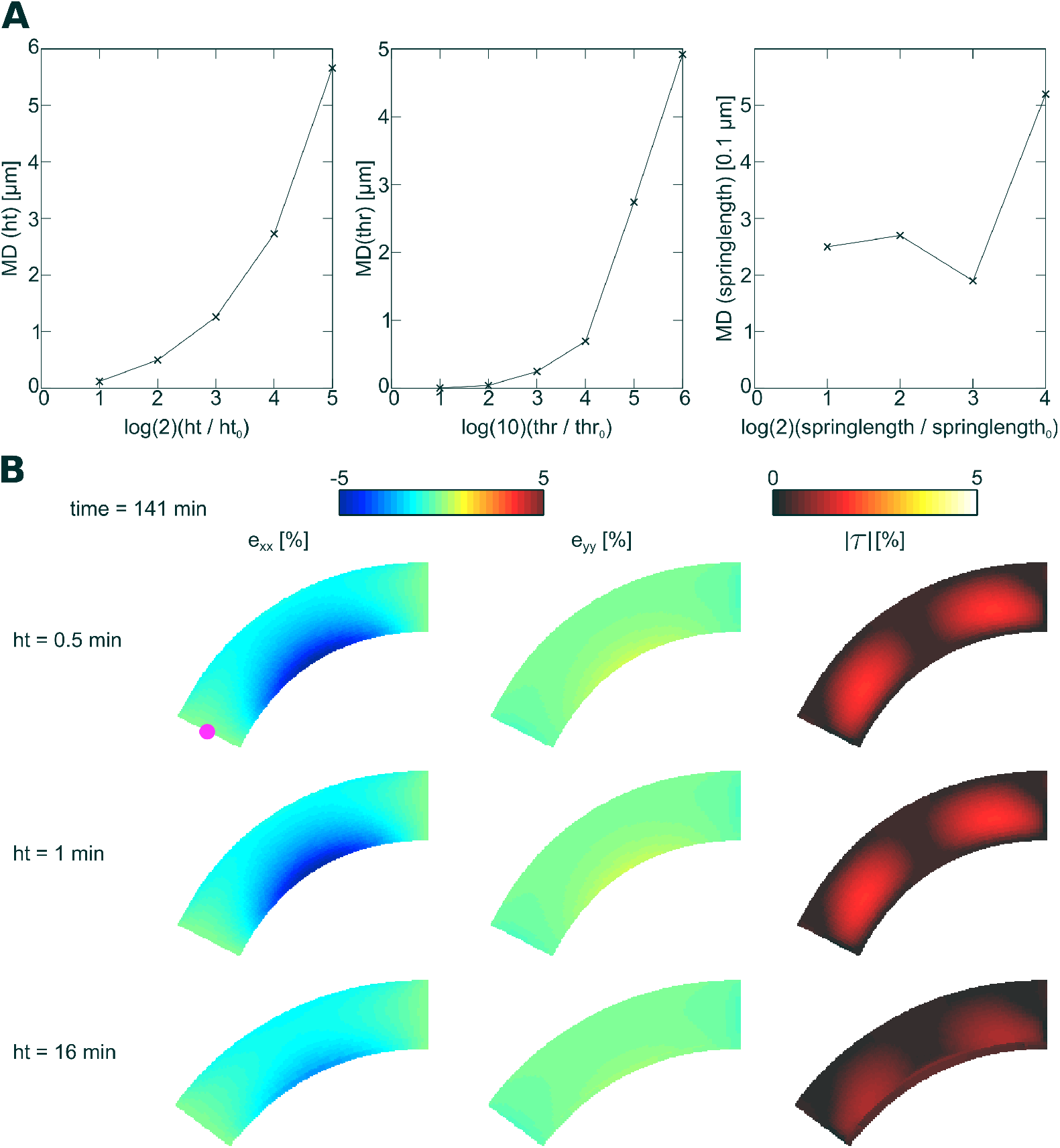
Validation of coupling parameters. **(A) Parameter setting causes convergent results**. Mean deviation (MD) of mass point trajectory as a function of coupling parameters. Reference trajectory was computed using strict parameter settings (*ht* = 0.5 *s*, springlength = 0.5 *μm*, *thr* = 0.005 *μN*). **(B) Emergent tissue bending and strain as a function of** *ht*. Mass point used for computing trajectories is illustrated by a violet point in left subfigure.

#### Damping constant

The equations for the motion of the mass points (Eqs. 18–26) are a system of coupled, damped, mechanical oscillators. We performed test simulations in which we used half and quartered values for the damping constant *η* to compute the viscous forces (Eq. 25). We found that these stricter settings do not affect the results, but only cause a slower convergence of the model. Thus, our parameter choice results in fast convergence of Eqs.26, with no numerical instabilities and convergent results.

#### Coupling parameters

To validate the settings for the parameters that couple the growth and mechanical models we performed the illustration experiment “anisotropic elongation” (setup is explained in detail in section Anisotropic elongation: “root bending”), and recorded a reference trajectory of a mass point located at the middle of the left side (“root tip”) of the model (see red point in left subfigure of Fig. 5B) using strict parameter settings (*ht* = 0.5 *s*, springlength = 0.5 *μm*, *thr* = 0.005 *μN*). Then we repeated the simulation with less strict parameter settings and computed the mean deviation of the trajectories relative to the reference trajectory. We computed the mean deviation (MD) for a parameter setting using

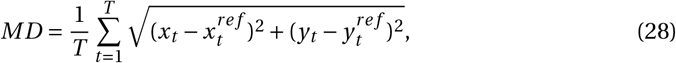

where 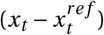 and 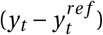 are the differences between the mass points positions on the x- and y-coordinate and the reference positions at time *t*. The trajectories were recorded for a simulation time of 141 *min*. Fig. 5A shows that trajectories are converging against the reference trajectory for increasingly strict parameter settings. We also see that our parameter choice leads to small mean deviations of the mass point trajectory (< 1 *μm*). The length of the reference trajectory is 123 *μm*.

How does the parameter setting affect the emergent strain fields during growth? To answer this question we illustrate these observables in Fig. 5B for the reference simulation (upper subfigure), our parameter choice (middle subfigure), and the parameter setting which caused the largest MD of ≈ 6 *μm* (bottom subfigure). We see that the reference simulation and the simulation with our parameter choice result in very similar results. The simulation with the least strict parameter setting results in a lower curvature of the tissue and lower residual strain; however, the qualitative results are identical. In conclusion we see that our parameter choice results in convergent results.

## Results

### Effect of finite strain on material properties

Here we characterize how finite strain and plastic growth affects the material properties of our model in numerical simulations. We start with illustrating the effect of finite strain on material properties in an isotropic setup without turgor pressure. We set elastic parameters in the model to *Y*_*x*_ = *Y*_*y*_ = 20 *MPa* · *m*; *v* = 0.1. Note that we use for efficient elastic properties (what we measure) a super- or sub-script “eff”, for instance *Y*_eff_ for the efficient Young’s modulus. We show the results in Fig 6. In Fig 6A,top we illustrate the setup of the direct stress experiment. We applied uniform, direct stress to the upper boundary of the model, while the bottom of the model was kept fixed on a horizontal line. Fig 6A, middle shows a stress-strain plot of the direct-stress experiment (black line) and as a comparison the theoretical behavior of a material which follows Hooke’s law (red line). We see that for small stress the model’s behavior converges to the theoretical value. For increasing stress however, the model stiffens, and a higher direct stress is required to stretch the model. Fig 6A, bottom shows the effective Young’s modulus and Poisson ratio against the strain (*ϵ*_*yy*_). We see that the effective Young’s modulus (solid line) increases linearly with increasing strain (with slope ≈ 2 *MPa* · *m*/10%), while the Poisson ratio maintains its theoretical value (dotted line). Note that with the highest direct strain value of 5% applied in the result section of this paper (compare Figs 10,11), the effective Young’s modulus is ≈ 10% larger than that for a theoretical isotropic Hooke’s material (without turgor). The linear increase in the stiffness of our simulated plant tissue is likely a result of our approximation to disregard the derivative of surface area A against strain (in Eq.8), which results in an error scaling linearly with direct strain (see Appendix **A**).

**Fig 6.**
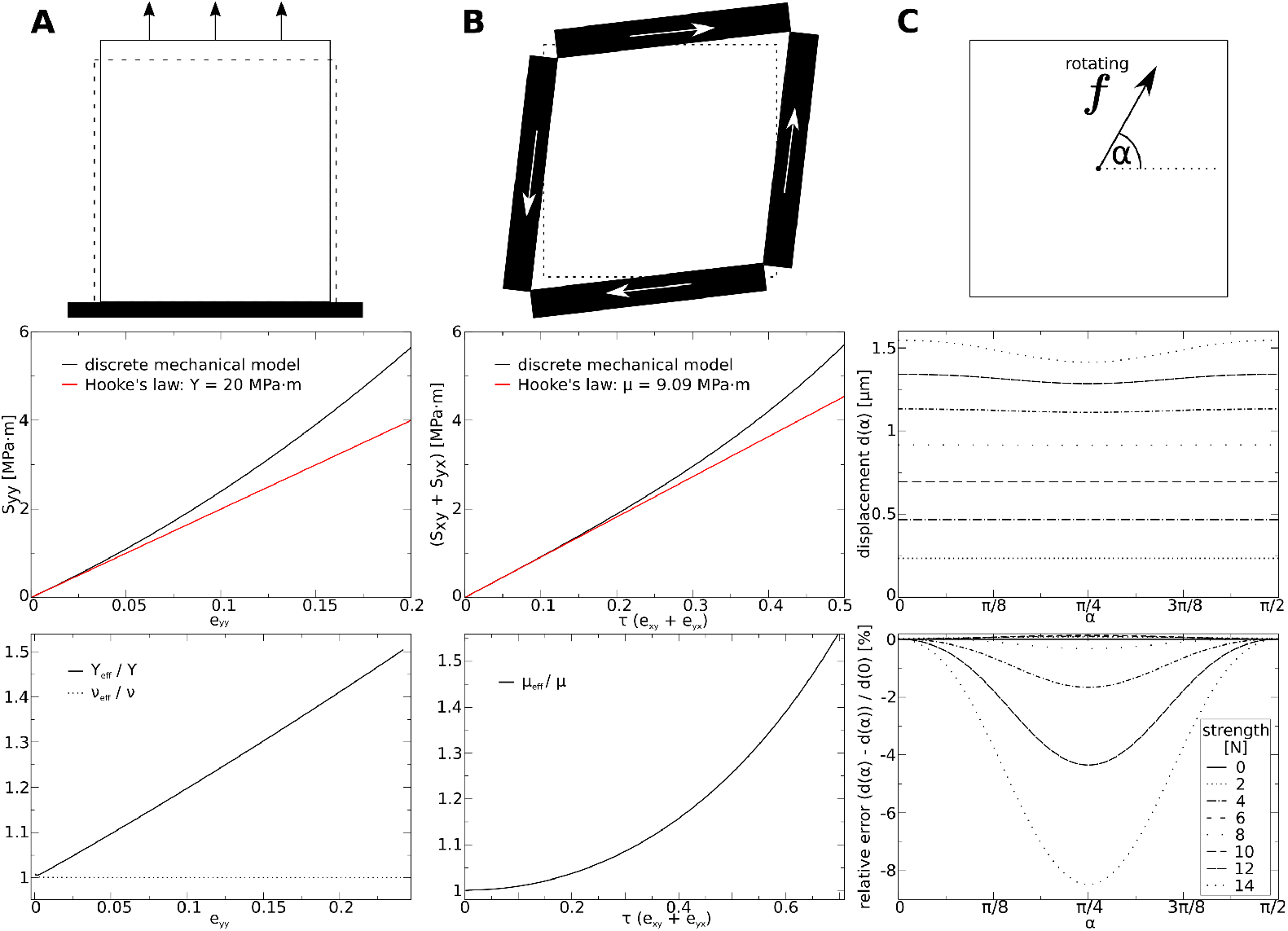
Illustration of material properties (isotropic, without turgor pressure). Top: Setups. Contours indicate undeformed (dotted) and deformed medium. Big arrows in **(A)** and **(B)** indicate applied stress. **(A) Direct stress experiment**. Top: Setup. Medium is fixed at the bottom (black block), such that points at lower border can only move horizontally. Middle: stress-strain plot. Bottom: ratio between the effective and theoretical values (*Y*_eff_/*Y*, *v*_eff_/*v*) as a function of *ϵ*_*yy*_. **(B) Pure shear stress experiment**. Top: Setup. Black blocks indicate walls through which shear stress is applied. Middle: stress-strain plot. Bottom: ratio between effective shear modulus *μ*_eff_ and theoretical *μ* as a function of *τ*. **(C) Isotropy experiment**. Top: Setup. Contour indicates fixed boundary of medium. Arrow symbolizes rotating force. Middle: displacement of center point for different strengths of force *vs* angle of force vector *α*. Length of undeformed squared medium 100 *μm*.

Next, we tested the shear properties of the model. In Fig 6B, top we illustrate the setup of this experiment. We applied shear stress *S*_*xy*_ and *S*_*yx*_ of same strength. Fig 6B, middle shows a stress-strain plot of the shear-stress experiment (black line) and as a comparison the theoretical behavior of a material which follows Hooke’s law (red line). We see that for smaller stress the model’s behavior converges to the theoretical value. For increasing stress however, again the model stiffens. Fig 6B, bottom shows the effective shear modulus *μ*_model_ against the strain (*ϵ*_*yy*_). We see that the effective shear modulus increases non-linearly with increasing strain. For a substantial total shear strain of *τ* = 10 % the effective shear modulus stifens 1 % compared to the theoretical value. Note that for the highest total shear strain of 5% which we chose as largest strain value in this paper, the model’s effective shear modulus is ≈ 0.3% larger than that following from the theoretical isotropic Hooke’s material (without turgor). This non-linear increase in the shear stiffness is likely also a result from our approximation to disregard the derivative of *A* with respect to strain. We showed in Appendix **A** that with respect to shear this approximation causes an error which scales quadratically with shear strain.

We also tested the isotropy of our model. We illustrate the experiment illustrated in Fig 6C. The rotating force, whose amplitude increases every rotation, was applied at the center point of a simulated tissue whose boundaries are fixed. To rule out boundary effects, we compared results of a medium of double the side length, and found qualitatively similar results. Fig 6C, middle depicts the displacement of the center point against the strength and the angle of the applied force. It reveals that that for forces smaller than 10 *N* the relative error of the mass point’s trajectory are smaller than 2%. However, it also shows that larger forces on the center mass point cause a substantial artificial anisotropy (error larger than 4%) in the model. A maximal strength of 14 *N* was applied in this study to the center point, which results in a substantial deformation gradient in the model. Note that this strength of force, if applied at all the boundary points, would correspond to a hydrostatic pressure of 14 *MPa* · *m*, which is one magnitude larger as the turgor pressure typically measured in *Arabidopsis thaliana* [56]. From Fig 6C, bottom we show the deviation from isotropic behavior against the strength and the angle of the applied force. We see that for forces smaller than 12 *N* the error is smaller than 4 %, and that for stronger forces the error substantially increases. In the simulations in the results section of this paper such localized strong forces do not occur, and thus we think that the model approximates the generalized Hooke’s law well.

### Effect of turgor pressure on material properties

We also studied the effect of finite strain on material properties in presence of turgor pressure. Before we present the simulation results, let us analyze what we expect to measure.

#### Analytical predictions

Shear modulus is defined (compare Eq. 2) as

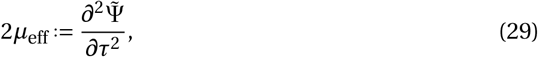

with *μ*_eff_ being the effective shear modulus (note difference to *μ*, the parameter). Taking into account the hydrostatic potential of a unit surface element in the elastic energy density (adding Eq. 16 for a unit cell to the energy density), we get

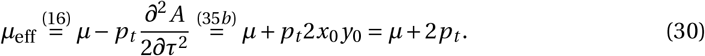

Note that *x*_0_ and *y*_0_ are unit lengths (*x*_0_ = *y*_0_ = 1). Thus, we expect to measure in simulations that the effective shear modulus is increased by twice the turgor pressure. Let us make a prediction based on the easiest example. If we disable the hinges in the model (set *μ* = 0), then we expect to measure as the model’s effective shear modulus *μ*_eff_ = 2 × *p*_*t*_.

Let’s now try to make predictions on how turgor pressure affects the effective Young’s modulus (*Y*_eff_) of the model (again: note the difference to the parameter *Y*). The effective Young’s modulus is defined as

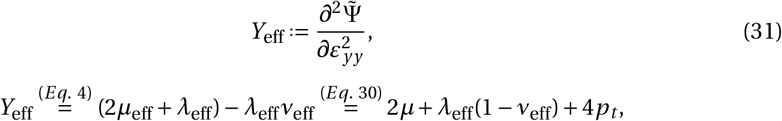

where *λ*_eff_ and *v*_eff_ are the effective Lame’ coefficients. Let’s assume that *v*_eff_ ≈ *v* and *λ*_eff_ ≈ *λ* (we will see later in simulations that this is reasonable). With this we get

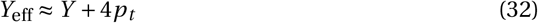

Thus we expect that the effective Young’s modulus of the model is approximately 4 × *p*_*t*_ stifer than without the added turgor pressure.

#### Numerical simulations

Now, let’s look at the simulation results, and compare it to our predictions. As described above, we first simplified the system, and performed shear experiments on the model without the hinges, thus setting parameter *μ* = 0, and making turgor pressure *p*_*t*_ solely responsible for resisting shearing forces. We measure in simulations for effective shear modulus due to turgor pressure: *μ*_eff_ = 2 × *p*_*t*_, for instance: (*p*_*t*_ = 1 *MPa* · *m*,(*S*_*xy*_ + *S*_*yx*_) = 0.1 *MPa* · *m*) resulted in *τ* = 5.00%, and thus *μ*_eff_ = (0.1/0.05)*MPa* · *m* = 2.00 *MPa* · *m*. This precisely fits our analytical arguments in Eq.30. Next, we performed the shear experiment in presence of turgor pressure (see Fig 7A). We see that, as expected from the above arguments, the model’s effective apparent shear modulus increases approximately linearly with increasing turgor pressure. However, we observe a larger stiffening effect than expected: for *p*_*t*_ = 0.25 *MPa* · *m* we see an increase of ≈ 0.6 *MPa* · *m* (instead of 0.5 *MPa* · *m*), and for *p*_*t*_ = 1 *MPa* · *m* we see an increase of ≈ 2.73 *MPa* · *m* (instead of 2.0 *MPa* · *m*). This is stiffening is probably due to the prestrain caused by turgor.

**Fig 7.**
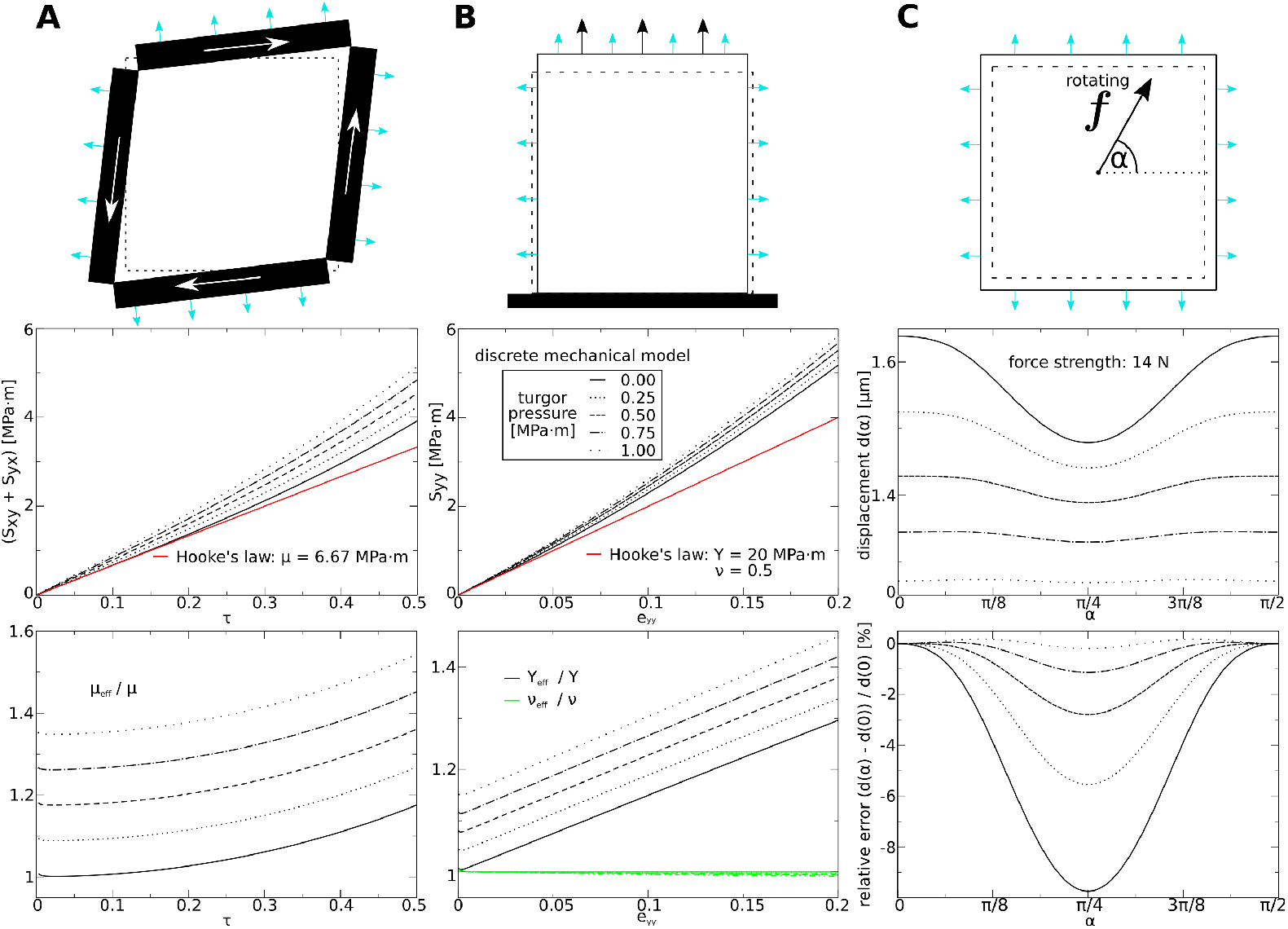
Material properties as function of turgor pressure. Top: Setups. Contours indicate undeformed (dotted) and deformed medium. Big arrows in **(A)** and **(B)** indicate applied stress. Blue arrows indicate turgor pressure. **(A) Shear stress experiment**. Top: Setup. Contours indicate undeformed (dotted) and deformed medium. Black blocks indicate walls through which shear stress is applied (all identical). Middle: stress-strain plots for different strengths of turgor pressure. Bottom: ratio between effective and theoretical shear modulus *μ*_eff_/*μ* as a function of *τ*. **(B) Direct stress experiment**. Top: Setup. Medium is fixed at the bottom (black block), such that points at lower border can move horizontally, but not vertically. Middle: stress-strain plots for different strengths of turgor pressure. ratio between effective and theoretical Young’s modulus and Poisson ratio (*Y*_eff_/*Y*, *v*_eff_/*v*) as a function of *ε*_*yy*_. **(C) Isotropy experiment**. Top: Setup. Arrow in the center symbolizes rotating force. Middle: displacement of center point for different turgor pressure and same strength of force (14 *μN*) *vs* angle of force vector *α*. Length of undeformed squared medium 100 *μm*.

In Fig 7B we show the results of the direct stress experiments in the presence of turgor pressure. We see that turgor pressure causes an increase in the model’s effective Young’s modulus. The effective Young’s modulus increases linearly with increasing turgor pressure, by ≈ 4 × *p*_*t*_ (see Fig 7A, bottom). For example we measure for a turgor pressure of 0.25 *MPa* · *m* and total strain of 1% an increase of the effective Young’s modulus 0.9 *MPa* · *m*. For a turgor pressure of 1.00 *MPa* · *m* and total strain of 4% we measure an increase of the effective Young’s modulus 3.5 *MPa* · *m*. Here we see a smaller stiffening effect than expected from our theoretical arguments. We see from Fig 7A, bottom that the effective Poisson’s ratio *v*_eff_ is affected very little by turgor pressure.

Finally, we studied how turgor pressure affects the model’s isotropy. The results are illustrated in Fig 7C. We added the turgor pressure to the medium first, fixed the boundaries of the medium, and then performed a similar experiment as in Fig 6C (but with different *p*_*t*_). Here we varied the strength of the turgor pressure, and kept the strength of the rotating force constant. Notably, we applied a strong rotating force of 14 *N* for which without turgor a substantial deviation from isotropy was observed. We can see from Fig 7C, middle that the displacement of the central mass point substantially decreases for higher turgor pressure. We can understand this from the results above, which demonstrated that the stiffness of the model increases with larger turgor pressure. In a stiffer medium we expect a smaller strain, and thus also a smaller deformation gradient.

### Effect of anisotropy on material properties

Here we show how anisotropy affects the material properties of our model. For this we performed simulations in a model with: *Y*_*x*_ = 20 *MPa* · *m*; *v* = 0.2; *p*_*t*_ = 0.5 *MPa* · *m*, and different *Y*_*y*_ = *c* · *Y*_*x*_, where we used *c* to control the fiber anisotropy. Results of these experiments are shown in Fig 8. In Fig 8A we show results of a direct stress experiment (similar to Fig 7A), in which we stretched the model in y-direction for different levels of anisotropy *c*. Fig 8A, middle shows that for smaller strain the model (black lines) converges to the theoretical value of a material which follows an orthotropic Hooke’s law (red lines). Whereas Fig 8, right reveals that the stiffening effect of the turgor pressure (compare Fig 7A, bottom) is decreased for larger *c*. This can partly be explained by the decreased prestrain (for higher *Y*_*y*_) due to the turgor pressure: for *c* = 1 prestrain is *e*_*yy*_ = 2%, for *c* = 2 prestrain is *e*_*yy*_ = 1%, and for *c* = 4 prestrain is *e*_*yy*_ = 0.4%. In Fig 8A we found that this causes an increase of the effective Young’s modulus in y-direction 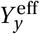: for *c* = 4 of 1%, for *c* = 2 of 2%, and for *c* = 1 of 4% Moreover, it is also shown that higher anisotropy *c* decreases the turgor pressure caused dependency on the effective Poisson ratio 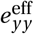. This may also be caused by the reduced stiffening due to prestrain in anisotropic conditions. So far, direct stress experiments were performed in only a single direction, as for isotropic tissues similar results would result from applying stress in the perpendicular direction. In contrast, for an anisotropic tissue model we need to apply stress experiments in x- and y-direction. In Fig 8B we show results of a direct stress experiment where stretch was applied in x-direction (compare Fig 8B, left). We see that the stress-strain plots coincide for different anisotropy ratios *c* (Fig 8B, middle). Consequently the plots 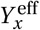 against *e*_*xx*_ (Fig 8B, right) are also very similar and again converge to the theoretical orthotropic Hooke’s law for small strains. We see also from (Fig 8B, right) that the effective Poisson ratio 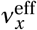 decreases with higher *s*. We can again explain this with the decrease in prestrain *e*_*yy*_ due to a higher stiffness of 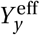 caused by anisotropy.

**Fig 8.**
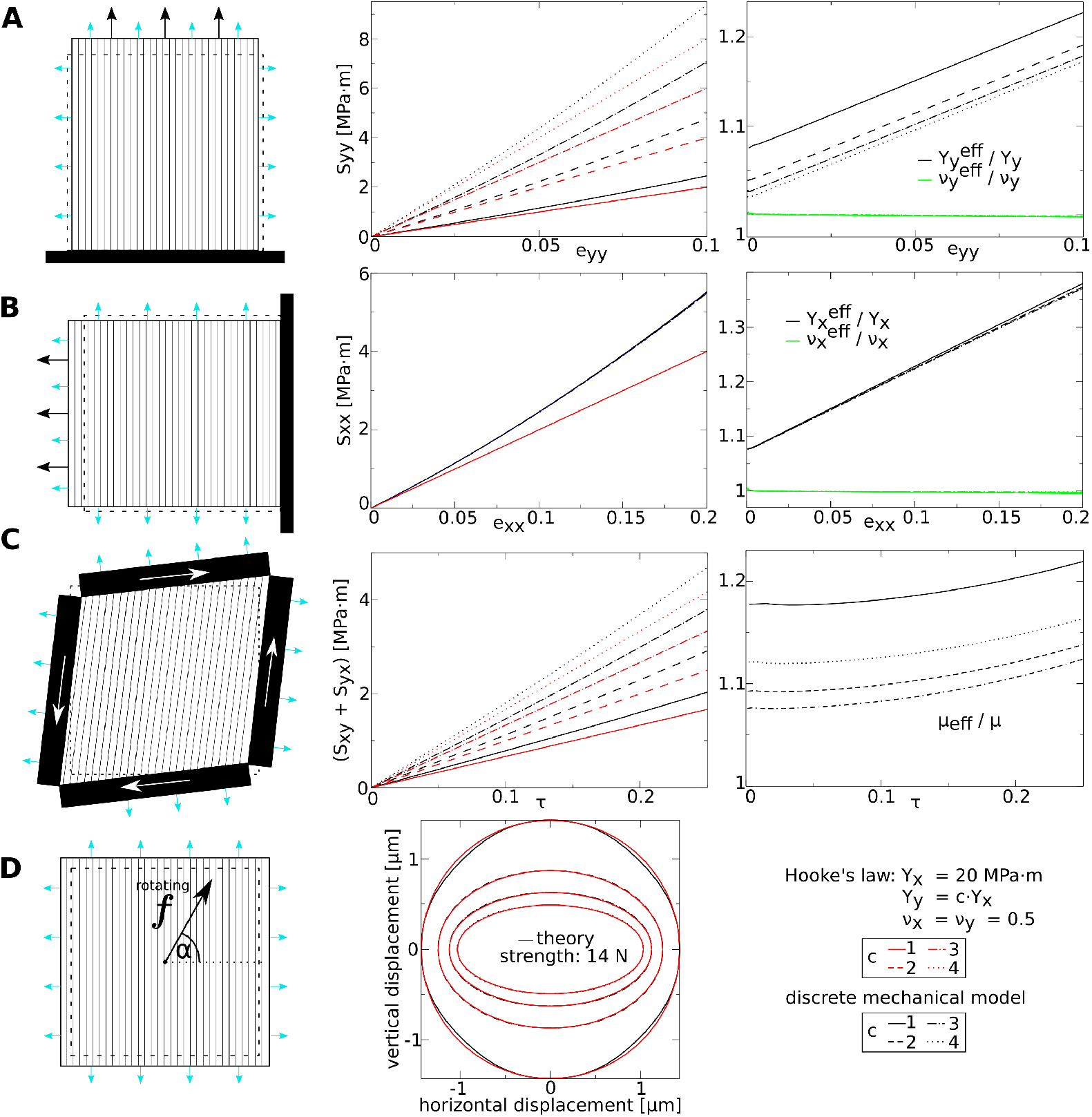
Material properties as a function of elastic anisotropy. Left: Fine lines indicate fiber direction, where *Y*_*y*_ = *c* × *Y*_*x*_. See Fig 7 for further explanation of setup illustrations. **(A) Direct stress experiment - stress in vertical direction**. Left: Setup. Middle: stress-strain plots for different anisotropy ratios *c*. Right: ratio between effective and theoretical Young’s modulus and Poisson ratio 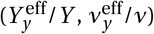 as a function of *ε*_*yy*_. **(B) Direct stress experiment - stress in horizontal direction**. similar to subfigure (A). **(C) Pure shear stress experiment**. Left: Setup. Middle: stress-strain plots for different anisotropy ratios *c*. Bottom: ratio between effective and theoretical shear modulus *μ*_eff_/*μ* as a function of *τ*. **(D) Isotropy experiment**. Top: Setup. Arrow symbolizes rotating force. Middle: displacement of center point for anisotropy ratios *c* and same strength of force (14 *μN*) *vs* angle of force vector *α*. Length of undeformed squared medium 100 *μm*. *p*_*t*_ = 0.5 *MPa* · *m*.

Next we performed the shear stress experiment on the model with different anisotropy *s* (Fig 8C). We see from the stress-strain plots in Fig 8C, middle that for small strain the effective shear modulus for different *c* (black lines) converges to the theoretical value of a orthotropic Hooke’s law (red lines). Additionally, we see for the shear modulus (Fig 8C, right) that a higher anisotropy *c* reduces the turgor pressure caused stiffening (compare to Fig 7B, bottom).

Finally we studied how well the anisotropy is described in the model by repeating the “rotating-force” experiment for a constant force strength of 14 *N* (compare Fig 7C) in models with different anisotropy ratios *c*. We plot the recorded trajectories (black lines) of the center point against theoretically expected ellipses (red lines). We constructed the ellipses using the initial position of the center point as the origin of the coordinate system, maximal displacement of the center point in x- and in y-direction (*x*_*p,max*_, *y*_*p,max*_) as vertices, and plotted for each *s* the ellipse:

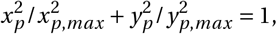

where *x*_*p*_ and *y*_*p*_ are the x- and y-coordinate of the center mass point. We see that the recorded trajectories match the assumed theoretical elliptic trajectories well.

### Effect of plastic growth on material properties

Above we demonstrated how the model’s material properties are affected by finite strain. Here we illustrate how plastic growth affects the material properties of the model. First similar experiments as shown above were performed, applying direct or shear stress while letting the medium grow uniformly. We show the results in Fig 9A-C. It can be seen from Fig 9A and B that the effective Young’s modulus *Y*_eff_ and Poisson ratio *v*_eff_ are affected little by plastic growth, for an increase from *h* = 1 (or *h*^−1^ = 1) to the maximal value of 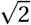 (when remeshing happens) changes in these material properties are less than 0.01%. However, from Fig 9C we can see that the effective shear modulus is affected substantially by plastic growth, it changes ≈ 6%, when *s* is increased from 1 to the maximal value of 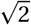. Moreover, the isotropy experiment (compare to Fig 6C) was repeated in a medium which we first let grow in horizontal direction. We show the relative error of trajectories of the center mass point (to which the rotating force is applied) for different values of *h*, and a strong local force of 10 *N* (compare Fig 6C). We find that the maximal change in metric of the unit cells due to growth causes a maximal relative error of ≈ 7% (compared to ideal circular trajectory).

**Fig 9.**
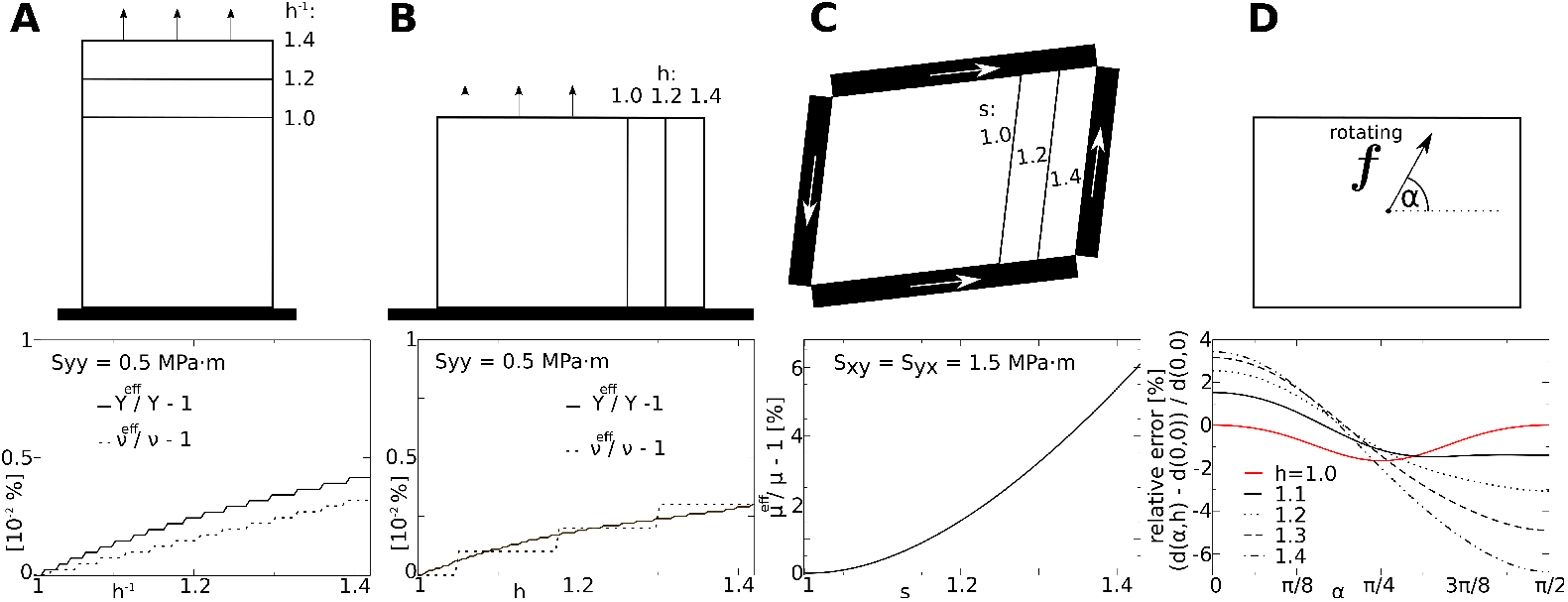
Material properties during growth. Fine lines indicate different system sizes (for different metric terms *h* and *s*). **(A) Direct stress experiment - growth in y-direction**. Top: Setup. Medium is fixed at the bottom (black block), such that points at lower border can move only horizontally. Black arrows indicate direct stress. Bottom: relative error of effective Young’s modulus and Poisson’s ratio as a function of *h*^−1^. **(B) Direct stress experiment - growth in x-direction**. similar to subfigure (A). **(C) Pure shear stress experiment**. Top: Setup. Black blocks indicate walls through which shear stress is applied. White arrows indicate shear stress (all same strength). Bottom: error of effective shear modulus as a function of *s*. **(D) Isotropy experiment**. Top: Setup. Contour indicates fixed boundary of medium (for *h* = 1.4). Arrow symbolizes rotating force. Bottom: error of trajectories for different *h* relative to displacement for *α* = 0, and *h* = 1.0. Length of initial (*h* = 1.0) squared medium 100 *μm*.

**Fig 10.**
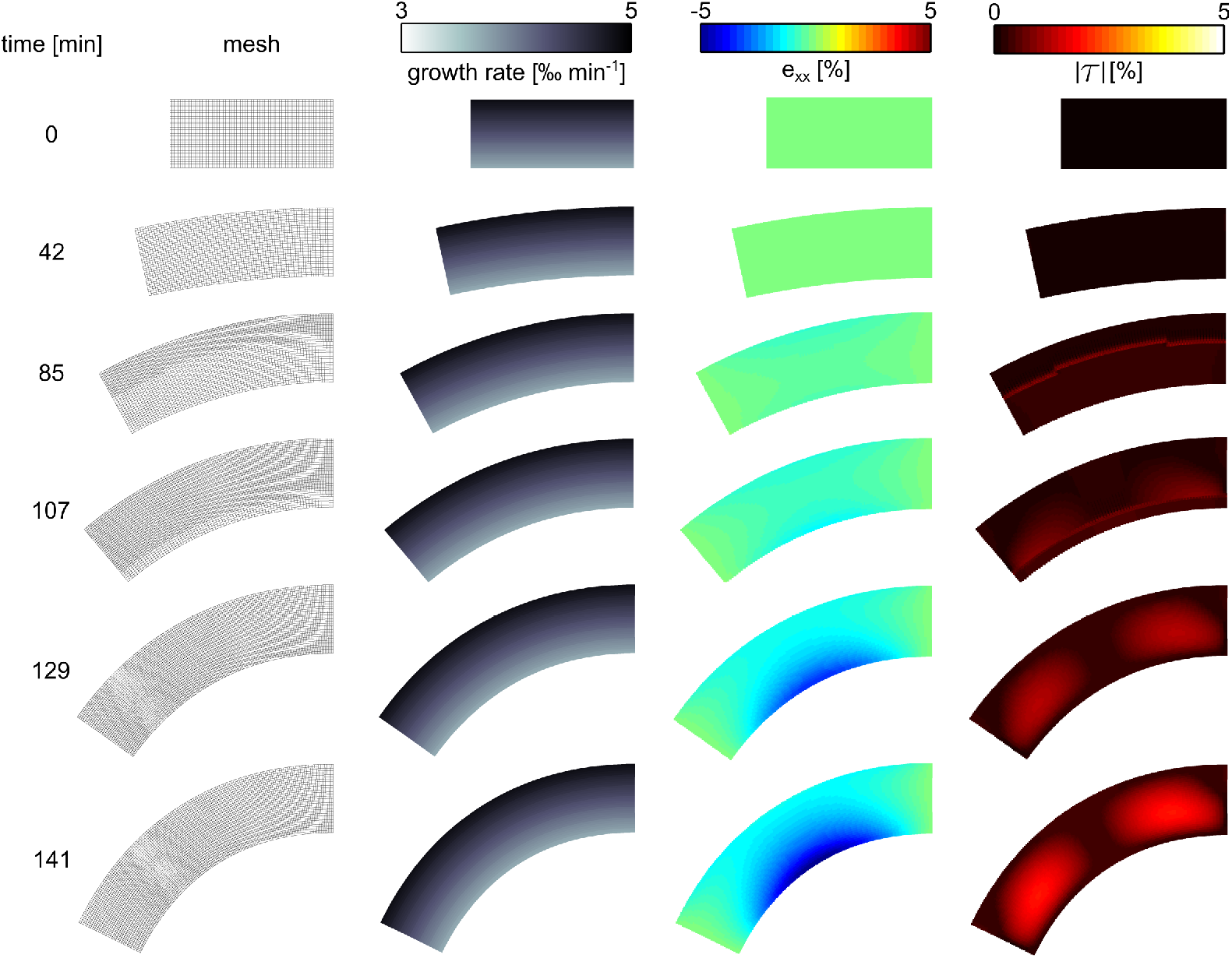
Application “root bending”. Anisotropic elongation (growth along root axis) causes emergence of residual strain and bending of the tissue.

**Fig 11.**
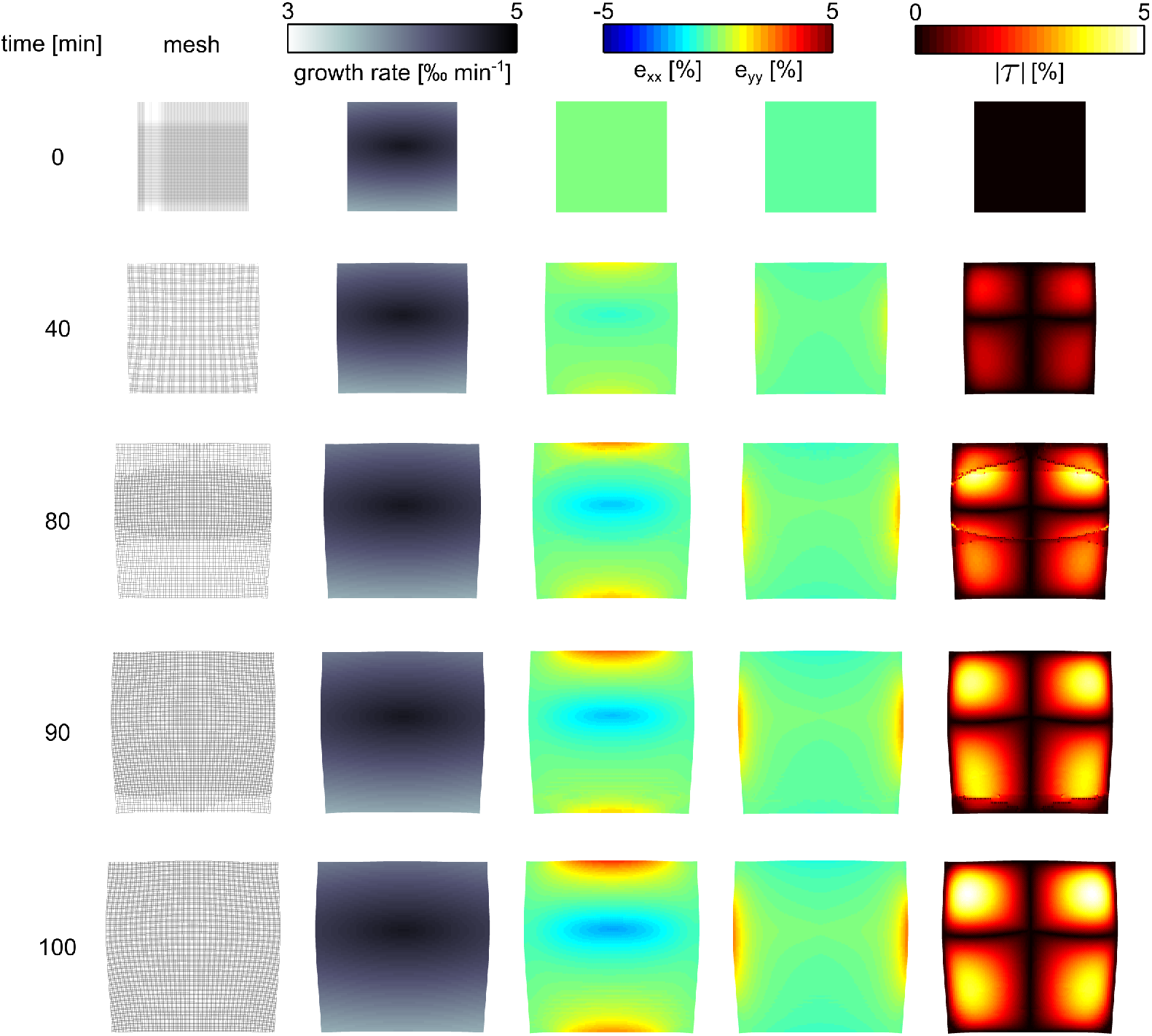
Application differential growth in 2D “leaf growth”. Anisotropic growth causes emergence of residual strain in the tissue.

We can conclude that growth causes a change in the metric of unit cells in our model, and that these changes affect the material properties mainly in its shear modulus. This artifact also manifests as a deviation from the ideal isotropic behavior. As a first application of our model we studied emergent residual strain fields and tissue deformation that arise due to anisotropic growth in rectangular model setups. We show data where a maximal residual strain of 5% emerges in the tissue, similar to previous modeling work [27].

### Anisotropic elongation: “root bending”

An important phenomenon in plant development is the bending of the root, for instance to grow towards nutrients, or to follow the gravitational field. Such directional growth responses are called tropisms, and they arise from a directional environmental signal becoming translated into a tissue level asymmetry of the plant hormone auxin. Since auxin levels dictate cellular expansion rates, this auxin asymmetry subsequently induces a growth rate asymmetry that results in bending. To illustrate the application of our method to the study of tropisms we superimposed a growth rate asymmetry.

To model root tropism, we use a slab of tissue with initial size 76 *μm* × 32 *μm*, with Young’s moduli *Y*_*x*_ = 40 *MPa* · *m* and *Y*_*y*_ = 80 *MPa* · *m*, Poisson’s ratio *v* = 0.2 and turgor pressure *p*_*t*_ = 0.2 *MPa* · *m*. We show the results of the simulation in Fig 10 and in video S1 Video. Growth happens in this setup only in x-direction (elongation along the “root axis”), and an asymmetric growth field is used (see Fig 10, second column) such that the upper part of the “root” grows faster than the lower part. We see that the asymmetric growth causes a bending of the tissue, where the inner side of the arc is the side with the slower growth rate. At 42 *min* the onset of bending can be seen. For time points 85 and 107 *min* a slight negative strain (compression) in the x-direction emerges (third column), which is weaker at the ends of the medium. In these snapshots we also see that remeshing is happening (left column) due to the growth process, starting from the outside arc, the location of highest growth rate and “propagating as a wave” towards the inner side of the arc. This remeshing does not cause a visible disruption of the direct strain in the x-direction (third column); however, for the absolute shear (right column) we see that the remeshing causes a slight local distortion of the shear strain field (see thin red line). These effects of the remeshing on the strain fields can be explained by our findings shown in Fig 9, where we showed that plastic growth affects the shear modulus, but hardly affects the Young’s modulus. Later the simulation (129, and 141 *min*) shows that the negative strain in the x-direction increases at the inner side of the arc (where the growth rate is smaller) to a maximum of −5%, whereas the strain on the fast growing side (outer side of the arc) is minimal. Furthermore we can see the emergence of a substantial shear strain field (see two “red eyes” in the right column). Note that throughout the simulation strain in the y-direction was very small, which is why we choose to not show it in Fig 10 (it is included in S1 Video).

### Anisotropic growth in two dimensions “leaf growth”

In many plant tissues, plastic growth is not restricted to a single direction. For example in leaf blades, tissue growth happens in two principal directions. Additionally, in case of bidirectional growth, tissue expansion is often anisotropic. Here we demonstrate such differential bidirectional growth in our model. For this we consider an initially quadratic slab of tissue with initial size 76 *μm* × 67 *μm*, with Young’s moduli *Y*_*x*_ = 40 *MPa* · *m* and *Y*_*y*_ = 80 *MPa* · *m*, Poisson’s ratio *v* = 0.2 and turgor pressure *p*_*t*_ = 0.2 *MPa* · *m*. We show the results of this simulation in Fig 11 and in S2 Video. We apply growth in x-, and y-direction, and use an asymmetric growth field (see Fig 11, second column). We see that residual strain fields emerge in the growing tissue. From Fig 11, third column we see that during growth positive strain in x-direction (stretch) gradually increases at the upper and lower border of the tissue, whereas a negative strain in the x-direction emerges in the center of the medium. In contrast to strain in the x-direction, in the y-direction weaker positive strain at the left and right borders gradually increases during growth, and no compression strain in the center happens. This can partly be explained by the elastic anisotropy of the tissue, where the stiffness in y-direction is twice larger compared to the x-direction (compare Fig 11, fourth column) causing that the tissue is easier deformed in x-direction than y-direction. We see that a substantial shear strain field (see four “red eyes” in right column) emerges in the medium.

## Discussion

We introduced a discrete mechanical growth model to study plant growth and development. The model contains an orthogonally organized mesh of mass points and connecting springs, providing an intuitive resemblance to the typical orthogonal microfibril architecture of anisotropic, polarized plant cells. We approximate the model’s material properties as an orthotropic Hookean material. The discrete nature of the method enables the incorporation of experimental data on material properties on the subcellular level. Compared to continuous mechanics approaches our method is relatively easy to implement in a computationally efficient manner, and allows usage of simpler integration schemes. We propose the method as a building block for multi-process models enabling researchers to link gene regulation, hormonal signaling, water transport and cellular behavior to the mechanics of tissue growth and deformation.

The model was used to study the consequences of growth. We found that anisotropic growth causes emergent strain fields in the medium, and that an asymmetric elongation (similar to root growth) causes a bending of the tissue. As a next step it is important to compare our predictions regarding tissue bending and emergence of strain fields to continuum mechanics models, and test them experimentally. Thusfar, the role of strain fields in root tropisms has not been investigated. Potentially the feedback of strain on growth mechanics could play a role in regulating growth asymmetry.

The model’s material properties were characterized in a series of simulations, and we discussed deviations from the approximated material law that arise from finite strain and plastic growth. A further limitation regarding material properties emerges from our choice of the coordinate system (fiber directions) to couple the discrete method to a continuous material. In presence of shear the underlying coordinate system is not ideally orthogonal but skewed. If under such conditions direct stress is applied, the “Poisson effect” in the model will artificially cause a force which is not perfectly orthogonal to the stress, but skewed. Importantly, for multi-scale biological models, the aim is to uncover how interactions between different processes shape tissue growth, development and adaptation rather than making precise predictions. In addition, typically, such research requires large numbers of exploratory simulations that probe different possible interactions between processes, initial conditions and parameter regimes, as opposed to computing a few distinct scenarios. Thus, we argue that the computational and numerical simplicity of our model outweighs its limitations in terms of accuracy for the aims it was developed for.

The model can be extended in various directions. For example, because of the discrete nature of the method breakage of cell wall material can be modeled with a removal of mass points, springs and hinges. This can be useful to model lateral root emergence when the forming tissue breaks through upper cell layers. In this paper we used simple growth functions to illustrate the model. However, as a next step growth should be formulated in terms of physiological processes. For instance, local growth can be formulated in terms of local auxin concentrations, strain fields and cytoskeletal processes. Moreover, the method may be applied not only to study plant tissue, but also to study other turgoid cell types of bacteria or fungi.

## APPENDIX

## A

In this section we demonstrate that for small strain, the following approximation used in Eq.8 is valid

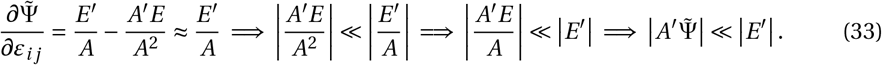

First, we write *A* in terms of strain

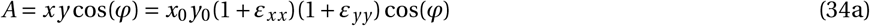

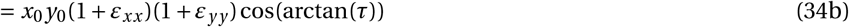

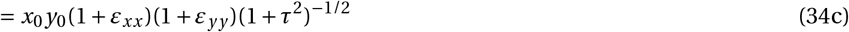

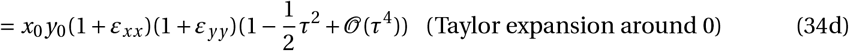

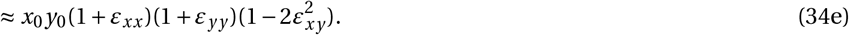

Taking the derivatives with respect to strain, and approximating these by keeping only terms that are of the lowest order in ***ε*** we obtain

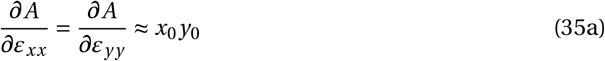

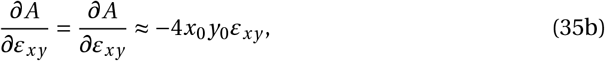

from this it follows that 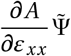 is of 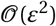, and 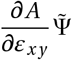 is of 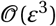,while *E*′ is linear with respect to strain components (compare Eq. 18). Therefore our approximation is valid for small strain.

Note that we can predict a scaling behavior of the errors for material properties for increasing strain. For the shear modulus *μ* we can expect an error to increase quadratically with increasing shear strain. Whereas we expect a linearly increasing error of the Young’s modulus *Y* for increasing direct strain.

## B

Here we explain the following approximation which was used in Eq.17

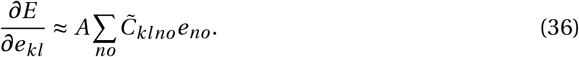

We use Eq.11 to write the left side of Eq.36 in terms of the stifness tensor

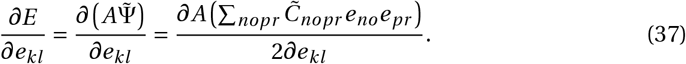

Because the elements of the stifness tensor *C*_*nopr*_ are proportional to *A* (see Eq.10), we define *H*_*nopr*_ := *C*_*nopr*_ /*A*, to rewrite the expression to

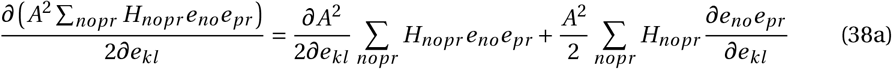

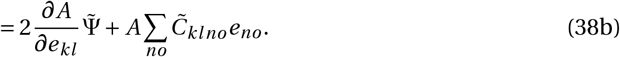

Thus our approximation (Eq.36) implies

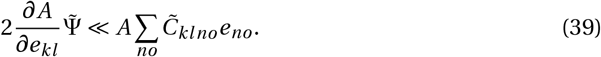

We again concentrate on terms with lowest order in ***ε*** (see Eq.35) to rewrite

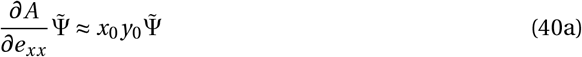

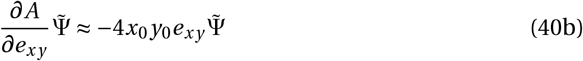

By taking the derivative with respect to direct strain components (*e*_*xx*_, *e*_*yy*_), we obtained expressions of 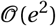, while by taking the derivative with respect to shear strain components (*e*_*xx*_, *e*_*yy*_) we obtained expressions of 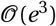. Finally we note that the right hand term in Eq.39 is linear with respect to strain components, and thus our approximation is valid for small strain.

## C

Here we derive Eq.22, the term which we use to compute the elastic shear force acting at a mass point

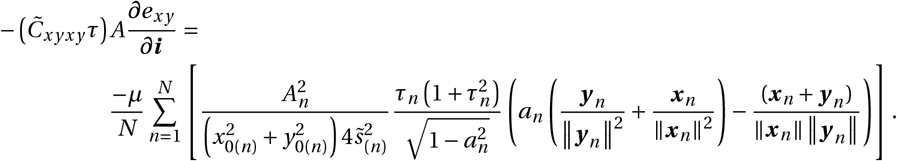

We start with the derivative 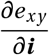:

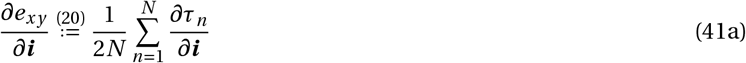

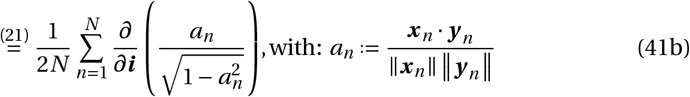

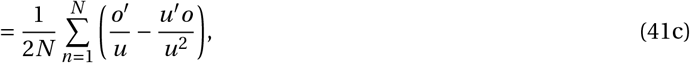

with:

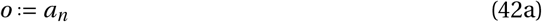

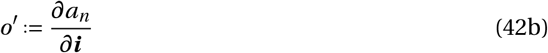

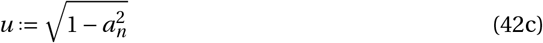

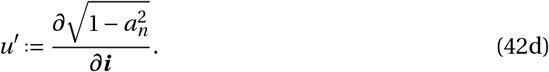

For the derivative 42b:

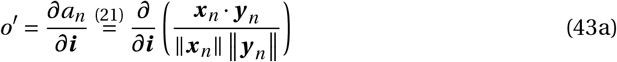

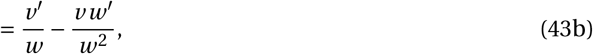

with:

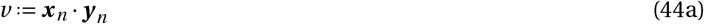

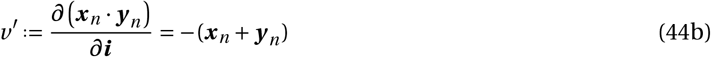

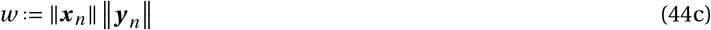

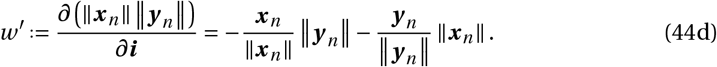

Substituting

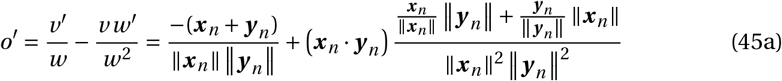

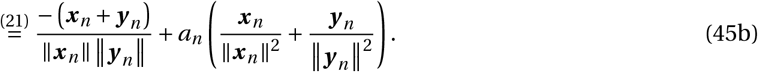

Considering the derivative 42d:

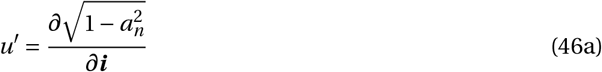

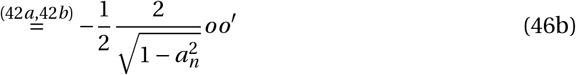

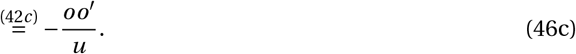

We rewrite Eq.41 into

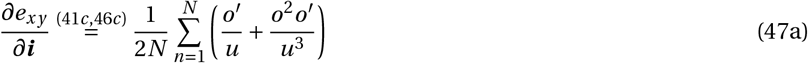

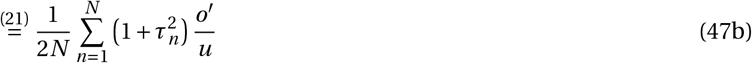

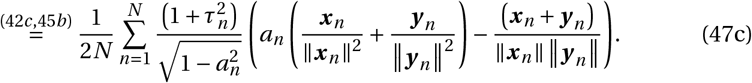

Finalizing the expression

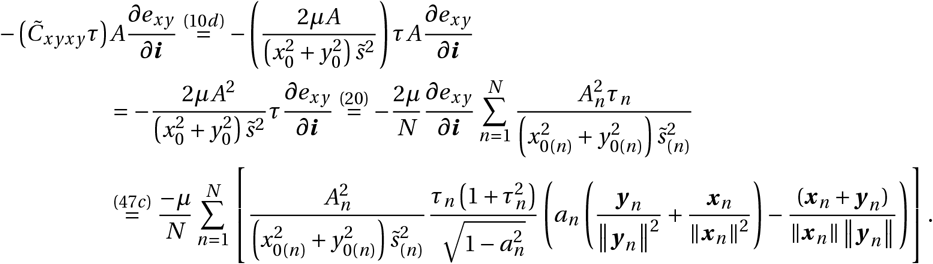

## Supporting information

**S1 File. Source Code.** The model was implemented using the C++ programming language. The software uses the Intel threading building blocks (TBB) runtime library as a parallelization environment (available in open source from http://www.intel.com/software/products/tbb/), and for visualization the CASH library from Rob J. de Boer and Alex D. Staritsky (available in open source from http://theory.bio.uu.nl/rdb/software.html).

**S1 Video. Application “root bending”.** Anisotropic elongation (growth along root axis) causes emergence of residual strain and bending of the tissue.

**S2 Video. Application “leaf growth”.** Anisotropic growth in x- and y-direction causes emergence of residual strain in the tissue.

## Acknowledgments

We thank Dr. Michael Sheinman, Sander Arens, Dr. Nadya Timofeeva, Dr. Matthias Mimault, Laurens Krah, and Dr. Chris van Dorp for valuable discussions and critical comments on the manuscript. We thank Dr. Paul Baron for help with language editing of the manuscript and Jan Kees van Amerongen for excellent technical support.

